# Separating measurement and expression models clarifies confusion in single-cell RNA sequencing analysis

**DOI:** 10.1101/2020.04.07.030007

**Authors:** Abhishek Sarkar, Matthew Stephens

## Abstract

The high proportion of zeros in typical scRNA-seq datasets has led to widespread but inconsistent use of terminology such as “dropout” and “missing data”. Here, we argue that much of this terminology is unhelpful and confusing, and outline simple ideas to help reduce confusion. These include: (1) observed scRNA-seq counts reflect both true gene expression levels and measurement error, and carefully distinguishing these contributions helps clarify thinking; and (2) method development should start with a Poisson measurement model, rather than more complex models, because it is simple and generally consistent with existing data. We outline how several existing methods can be viewed within this framework and highlight how these methods differ in their assumptions about expression variation. We also illustrate how our perspective helps address questions of biological interest, such as whether mRNA expression levels are multimodal among cells.

## Introduction

Single-cell RNA sequencing (scRNA-seq) has facilitated investigation of important biological questions that were previously difficult or impossible to study, such as the nature of heterogeneity within classical cell types, the dynamics of cellular processes, and the biological pathways underlying cellular differentiation. However, how to model scRNA-seq data has been the subject of considerable confusion and debate. In particular, the high proportion of zeros in typical data sets has garnered special attention, and has led to widespread but inconsistent use of terminology such as “dropout” and “missing data.” In this paper, we argue that much of this terminology is confusing and unnecessary, and we outline simple ways of thinking and talking about scRNA-seq data that can help reduce confusion.

The first key idea is that observed scRNA-seq counts reflect two distinct factors: the variation in actual expression levels among cells, and the imperfect measurement process. Therefore, models for observed scRNA-seq counts, which we will call *observation models*, are obtained by specifying: (1) an *expression model* that describes how the true expression levels vary among cells/genes, and (2) a *measurement model* that describes how observed counts deviate from the true expression levels. Distinguishing between observation, expression, and measurement models is important both for avoiding confusion and for performing useful analyses. Indeed, the goal of most RNA-seq analyses is to draw inferences about true expression levels from observed counts, and this is impossible without explicit consideration of how the observed counts are related to the expression levels through a measurement process. Moreover, making measurement and expression models explicit can help clarify the underlying assumptions and aid interpretation of results such as parameter estimates.

The second key idea is that a Poisson model is a reasonable starting point for modeling scRNA-seq measurement. We summarize theoretical arguments for this model, and explain how it can capture the abundance of zeros in scRNA-seq data without special terminology or special treatment. This measurement model is a simplification (Box 1), but we argue that it is a useful simplification that will often suffice in practice.

Both ideas are simple, and neither is new. Modeling the measurement process has a long history^1^, as does the use of Poisson measurement models for RNA-seq^2,3^. However, many papers on scRNA-seq analysis do not incorporate these ideas, focusing exclusively on observation models and leaving measurement and expression models implicit (ref.^4^ is a notable exception). This ambiguity is especially problematic when the models include components said to capture “zero-inflation” without clearly indicating whether these are part of the measurement model, the expression model, or both. Here, we show how many scRNA-seq observation models can be interpreted as combining a Poisson measurement model with different expression models, clarifying their underlying assumptions about expression variation.

These simple ideas can also help address questions of biological interest. For example, the question of whether gene expression patterns are multimodal among cells is about the expression model, not the observation model. We investigate this question empirically in diverse datasets, and find that data are often consistent with surprisingly simple expression models. Specifically, a Gamma distribution often suffices to capture variation in expression levels among cells.

## A call to simplify terminology

One major source of confusion in scRNA-seq analysis is the widespread but inconsistent use of terminology, especially “dropout,” “missing data,” “imputation,” and “zero-inflation.” Choice of terminology has many important consequences: it affects the way that researchers think, develop and apply methods, and interpret results. We therefore begin by reviewing these terms, and explain why in many cases we view them as unhelpful.

The term “dropout” has become commonly used in connection with the zeros in scRNA-seq data^5–9^. Historically, dropout referred to *allelic dropout*, a failure of PCR in which specific primers would fail to amplify sequences containing a specific allele, leading to genotyping errors for heterozygous individuals^10,11^. In scRNA-seq, the term “dropout” was introduced to describe a supposed failure that might cause a gene to appear highly expressed in one cell but not expressed in another^5^. Although the source was not specified, this usage seems to refer to some aspect of the measurement process. The term has since spread widely, but its meaning varies among papers and presentations (and sometimes even within papers and presentations!). For example, it is used to refer sometimes to observed zeros, sometimes to the unknown subset of zeros at genes that are expressed but undetected, and sometimes to the fact that not all molecules present in the original sample are observed (which affects all observations, not just the zeros). Such variation in usage naturally leads to confusion and disagreement about how these phenomena should be modeled. For this reason, we argue the term “dropout” should be avoided in the context of scRNA-seq data. Instead, zero observations should simply be referred to as zeros, and measurement models should focus on the fact that not all molecules in the original sample are observed, with zeros being just one byproduct.

The term “missing data” is also commonly used in connection with zeros in scRNA-seq^8,12^. This terminology is misleading because the zeros are not missing data as understood in statistics. To illustrate, consider a survey where a researcher counts the number of cars arriving at an intersection each minute. If no cars arrive in a particular minute, then the observation would be recorded as zero; however, this observation is not “missing.” In contrast, if the researcher takes a lunch break and does not record arrivals for an hour, then this would lead to sixty truly missing observations. The zeros in an scRNA-seq data experiment are more like the former than the latter: there is no analogue of a lunch break in the scRNA-seq measurement process. Of course, in RNA-seq data the observed zeros are noisy, and do not necessarily imply that there were zero molecules present in the original cell; however, the same is true of all observations.

Inappropriate use of the term “missing data” has led to inappropriate application of methods for dealing with missing data to scRNA-seq data. For example, in other applications it is common to “impute” (fill in) missing values, and so many scRNA-seq papers have described methods to “impute” zeros^8,13–15^. However, since zeros in scRNA-seq are not actually missing data, the meaning of these imputed values is unclear. Thus, the term “missing data”, and methods that “impute” (only) zeros, should be avoided. Instead, every observation in scRNA-seq should be treated as a noisy observation of an underlying true expression level, and inference should focus on clearly-defined tasks such as estimating these true expression levels.

Finally, the term “zero-inflated” is another common source of confusion and debate. In statistics, “zero-inflated” describes a model for count data that is obtained from a simpler model by increasing the proportion of zeros. For example, “zero-inflated Poisson” refers to a distribution obtained by taking a Poisson distribution and then increasing the proportion of zeros. In scRNA-seq applications, the use of zero-inflated models is an understandable reaction to the high proportion of observed zeros. However, recent work suggests that zeroinflated models may be unnecessary; indeed ref.^16^ entitles their piece “Droplet scRNA-seq data is not zero-inflated.” Here, we take a slightly different perspective: we argue that there is no convincing evidence supporting zero-inflated measurement models; however, zero-inflated observation models could be appropriate, depending on actual expression variation. This perspective makes clear that the need for zero-inflated models may vary among data sets and among genes.

These issues all stem from one major misconception in scRNA-seq: the idea that the measurement process involves some distinct zero-producing technical mechanism. Indeed, some published observation models include a component that randomly creates zero observations irrespective of the true expression level, for example ref.^14^. Such a mechanism would require a systematic effect that somehow misses all molecules from a particular gene in a particular cell, and there are no convincing theoretical arguments or empirical evidence supporting this idea. Thus, we argue that measurement models for scRNA-seq should begin from a simple assumption, that the measurement process operates independently on each molecule in the cell. This assumption is a simplification (Box 1), but is not strongly contradicted by existing data, and leads to simple observation models and explanations for the large proportion of zeros in scRNA-seq data.

## Modeling scRNA-seq data

Observed scRNA-seq counts reflect both the true expression levels of each gene in each cell, and the measurement process. We first describe measurement and expression models, and how they combine to yield observation models. We focus on data generated using Unique Molecular Identifiers (UMIs), which substantially reduce unwanted variation, including differences in gene lengths and PCR amplification efficiencies^17^. (It is possible that zero-inflated observation models were initially motivated by a need to account for variation in read counts introduced by PCR.) We ignore these sources of variation, so our arguments may not apply to data generated without UMIs.

We now introduce some notation. Consider using scRNA-seq to measure gene expression in *n* single cells. Thinking of each cell as a pool of mRNA molecules, let *m_ij_* denote the true (but unknown and unobserved) number of molecules present in cell *i* from gene *j* (*i* = 1, …, *n*;*j* = 1, … , *p*), and let *m_i+_* denote the total number of molecules in cell *i*. We refer to *m_i+_* as the *absolute expression level* of gene *j* in cell *i* and 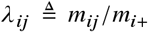 as the *relative expression level* of gene *j* in cell *i*. In words, the absolute expression level of a gene is the number of RNA molecules present from that gene, whereas the relative expression level is the proportion of RNA molecules from that gene. Here, we focus on relative expression levels since estimating absolute expression levels from scRNA-seq is difficult, and use **Δ** = [*λ_ij_*] to denote the matrix of relative expression levels. Let **X** = [*x_ij_* denote the observed count matrix, where *x_ij_* denotes the number of distinct molecules from gene *j* observed in cell *i*, and let *x_i+_* denote the total number of molecules observed in cell *i*.

In this notation, a measurement model is a model that connects the observed counts **X** to the expression levels **Δ**, by specifying the conditional distribution *p*(**X / Δ**). An expression model is a model for the expression levels *p*(**Δ**). Together, these two models determine the observation model, which is a model for the observed counts *p*(**X**).

### Modeling scRNA-seq measurement

We believe that methods should generally start with simple models, adding additional complications only when warranted. In this spirit, we suggest that methods for scRNA-seq should start with a simple Poisson measurement model:

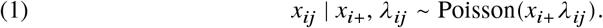

Although simple, this measurement model is supported by theoretical arguments^3,18^ (Supplementary Note 1), early empirical analyses of bulk RNA-seq data2, more recent analyses of control scRNA-seq data^4,16,18^, and our own empirical analyses.

Although the Poisson measurement model is essentially the same as that used for bulk RNA-seq data, it can nonetheless account for the fact that there are many more zeros in scRNA-seq data than bulk RNA-seq data. First, the total number of molecules observed *x_i+_* is typically much smaller for single cells than for bulk samples, because single cells have less starting material and are typically sequenced to lower average depth. Second, it is more common that *λ_ij_* will be small (or even zero) for a single cell than for a bulk sample. This is because expression levels in bulk samples are averages of expression levels in many single cells, and averaging reduces the frequency of both small and large values. These two facts imply that the rate parameter *x_i+_ λ_ij_* is often smaller in scRNA-seq data than in bulk RNA-seq data, explaining the higher proportion of zero observations.

The Poisson measurement model also captures aspects of scRNA-seq data that have previously been referred to using terms such as “dropout,” “missing data,” and “technical zeros.” It captures the fact that not every molecule that was present in every cell was observed; indeed, this fact is a fundamental assumption. It also captures the fact that *x_ij_* may be observed to be zero even when *m_ij_* (hence *λ_ij_*) is non-zero. However, it captures these features without introducing a distinct zero-generating mechanism. The zeros, like other observations, are simply imperfect measurements with no need for special terminology or special treatment.

Under the Poisson measurement model, observing *x_ij_* = 0 is different from *x_ij_* being missing. If *x_ij_* were missing, then it would provide no information about the expression level *λ_ij_*. But, observing *x_ij_* = 0 does provide such information, namely that *λ_ij_* is unlikely to be large. In other words, it correctly reflects that counts *x_ij_* are noisy observations of the true expression levels; see also refs.^19–21^ for example.

#### Alternative measurement models

Previous papers have considered zero-inflated measurement models for scRNA-seq data^5,8^. However, there are no convincing empirical analyses supporting such models; neither are there any convincing arguments for why such zero-inflation should be expected. Furthermore, two recent analyses of control data sets, in which synthetic mRNA molecules are directly added (“spiked in”) to droplets at known concentrations, captured, and then sequenced found no evidence supporting a zero-inflated measurement model^4,16^. Our empirical analysis below further supports this position. Including an unnecessary zero-generating component in the measurement process has the cost of increasing model complexity, and introduces the danger that true expression variation may be wrongly attributed to the measurement process.

This said, the Poisson measurement model (1) is necessarily a simplification. In particular, it ignores biases that may cause some molecules to be more likely to be observed than others (Box 1; Supplementary Note 1). When such biases are consistent across cells, inferences that involve comparisons among cells will be robust to ignoring them; however, some biases may vary from cell to cell, producing measurements that are overdispersed (more variable) relative to a Poisson distribution. Given the difficulty of precisely modeling all aspects of the measurement process, and given that available data do not strongly contradict a Poisson measurement model, our perspective is that methods development should start with the Poisson measurement model, and focus more attention on modeling expression variation (as described below). The main downside of this approach is that it risks overstating expression variation if measurement overdispersion is high. However, we view this as a risk worth paying for the benefits of simplicity.

### Modeling gene expression

We emphasize that our suggestion to use a Poisson model applies specifically to the measurement model, not the observation model. Indeed, many papers have demonstrated that a Poisson observation model does not capture all variation in observed RNA-seq data, and so it is common to use a more flexible observation model that can capture additional variation, such as NB or zero-inflated negative binomial (ZINB) observation models. These observation models are not inconsistent with a Poisson measurement model; indeed, in this section we explain how NB and ZINB observation models, as well as many other existing methods, naturally arise by combining the Poisson measurement model with certain expression models.

To make this idea precise, first consider modeling expression at a single gene *j*. An expression model for a single gene involves specifying a distribution *g_j_* for the expression levels *λ_1j_*, … , *λ_nj_*,

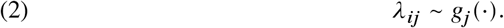

One simple choice is to assume that *g_j_* is a Gamma distribution; combining this with (1) yields the NB observation model^22^. Similarly, combining a *point-Gamma* expression model, wherein some proportion of the expression levels are exactly zero, while the other (non-zero) levels follow a Gamma distribution, with (1) yields the ZINB observation model. It is also possible to use non-parametric expression models^4,23^. A list of single-gene expression models, and some published methods implementing statistical inference for the corresponding observation models, are given in Table 1 (see Supplementary Note 2 for details).

**TABLE 1.** Single gene models for scRNA-seq data. Different expression models, when combined with the Poisson measurement model, yield different observation models. Method indicates previously published methods and software packages that use the corresponding observation model to analyze data.

| Expression model | Observation model | Method |
| --- | --- | --- |
| Point mass (no variation) | Poisson | Analytic |
| Gamma | Negative Binomial | MASS<sup>40</sup>, edgeR<sup>41</sup>, DESeq2<sup>42</sup>, BASICS<sup>43</sup>, SAVER<sup>19</sup> |
| Point-Gamma | Zero-inflated Negative Binomial | PSCL<sup>44</sup> |
| Unimodal (non-parametric) | Unimodal | ashr<sup>23,45</sup> |
| Point-exponential family | Flexible | DESCEND<sup>4</sup> |
| Fully non-parametric<sup>46</sup> | Flexible | ashr |

Similar ideas apply to expression models for multiple genes, although things inevitably become more complex. A multi-gene expression model simultaneously describes correlations among expression levels at different genes across cells, and different cells across genes. A common and powerful approach to describe these correlations is to use low rank models, which intuitively assume that the correlations can be captured by a relatively small number of patterns, much smaller than the number of cells or genes. More precisely, these models can be written

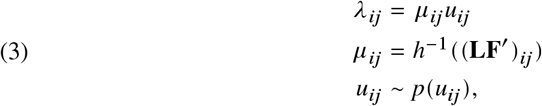

where *h* denotes a *link function*, or transformation, and the *loadings* matrix **L** and the *factors* matrix **F** are low rank. This form has an appealing biological interpretation: the matrix **LF’** represents the structure of expression variation among cells/genes and *u_ij_* represents stochastic deviations from this structure. Thus, one might think of **LF’** as representing different cell types/states and *u_ij_* as stochastic expression noise^24^. A list of multi-gene expression models, and some published methods implementing statistical inference for the corresponding observation models, are given in Table 2 (see Supplementary Note 3 for details).

**TABLE 2.** Multi-gene models for scRNA-seq data. Multi-gene models partition variation in true expression into structured and stochastic components. The link function describes a transformation, and the noise distribution indicates an assumption about the stochastic component (*p(u_ij_*) in (3)). Method indicates previously published methods and software packages that use the corresponding observation model to analyze data.

| Link function | Noise distribution | Method |
| --- | --- | --- |
| Identity | None | NMF<sup>47</sup>, scHPF<sup>48</sup> |
| Identity | Gamma | NBMF<sup>49</sup> |
| log | None | GLM-PCA<sup>18</sup> |
| log | Gamma | scNBMF<sup>50</sup>, GLM-PCA<sup>18</sup> |
| log | Point-Gamma | ZINB-WaVE<sup>51</sup> |
| Neural network | Point-Gamma | scVI<sup>28</sup>, DCA<sup>20</sup> |

Interestingly, methods that combine expression models of the form (3) with the Poisson measurement model (1) may be robust to misspecification of the measurement model. Specifically, if measurements are overdispersed relative to Poisson, then the model fit will tend to include this additional variation in *p(u_ij_*) (provided this distribution is sufficiently flexible), while leaving estimates of the structured variation **LF’** unchanged. Intuitively, overdispersered measurement error affects the variance of the observations, but not the mean. Therefore, while estimates of the stochastic noise may be sensitive to assumptions on the measurement process, estimates of the structured expression variation will be more robust.

### Summary

To summarize, many existing observation models for scRNA-seq data can be derived by combining the Poisson measurement model (1) with expression models of the form (2) or (3). This framework clarifies several sources of confusion in scRNA-seq analysis. First, it emphasizes that which observation models are most appropriate for scRNA-seq data may vary among data sets and among genes, because expression variation will vary among data sets and genes. For example, if data are collected on a set of homogeneous cells, then most genes might show relatively little expression variation may be adequately described by simple expression models. In contrast, if the data contain many cell types, then more complex expression models could be required. We study this question empirically below.

Second, this framework provides a different interpretation of a finding, for example, that a ZINB observation model fit observations at some gene in some data set better than an NB model: it would imply that the expression levels at the gene are better modeled by a point-Gamma distribution than by a simple Gamma distribution. Importantly, this is a conclusion about the true expression levels, not a conclusion about the measurement process. This interpretation contrasts with the usual way that the ZINB observation model is interpreted, in which zero-inflation captures some supposed technical mechanism (see Supplementary Note 4 for details).

Finally, this framework provides a rigorous approach to infer, for example, the mean or variance of true gene expression levels, as well as the true expression levels themselves, from the observed counts. However, these estimates are generally neither simple functions of the observed counts, nor functions of simple transformations (e.g., log) of the observed counts. We illustrate this procedure for a single-gene point-Gamma expression model (ZINB observation model) in Box 2.

## Empirical examples

### Single gene models

There is considerable debate about whether scRNA-seq data are adequately modeled by an NB observation model, or if it is necessary to use a ZINB observation model. Some papers have concluded that observed scRNA-seq data exhibit multi-modal expression variation, suggesting that an even more complex observation model may be necessary^25–27^. Under the framework outlined above, these questions translate into questions about the expression model: is a Gamma expression model adequate, or is it necessary to use a more complex, even multi-modal, expression model?

Since expression variation may vary among genes and data sets, we analyzed data sets from a range of settings including homogeneous collections of sorted cells, *a priori* homogeneous cell lines, and heterogeneous tissues. We also created *in silico* mixtures of sorted cells as positive controls for highly heterogeneous expression patterns (Table 3).

**TABLE 3.** Data sets analyzed. Number of samples passing the previously reported QC filters and number of genes with non-zero observations in at least 1% of samples, or passing the previously reported QC filters53. (a) Mixture data sets are generated *in silico* by concatenating the data and then applying QC filters. (b) Data downloaded from https://10xgenomics.com/data.

| Dataset | Protocol | Number of samples | Number of genes | Source |
| --- | --- | --- | --- | --- |
| (Sorted) T cells | GemCode | 10,209 | 6,530 | <sup>52</sup> |
| (Sorted) B cells | GemCode | 10,085 | 6,417 | <sup>52</sup> |
| iPSC | Fluidigm C1 | 5,597 | 9,957 | <sup>53</sup> |
| T cell/B cell mix<sup>a</sup> | GemCode | 20,294 | 6,647 | <sup>52</sup> |
| Cytotoxic T/Naive T mix<sup>a</sup> | GemCode | 20,688 | 6,246 | <sup>52</sup> |
| Brain | DroNc-Seq | 14,963 | 11,744 | <sup>54</sup> |
| Kidney | 10X Chromium v2 | 11,233 | 15,496 | <sup>55</sup> |
| PBMC | 10X Chromium v3 | 11,769 | 12,144 | <sup>b</sup> |
| Retina | 10X Chromium v2 | 21,285 | 10,047 | <sup>56</sup> |
| Control 1 | 10X Chromium v2 | 2,000 | 88 | <sup>57</sup> |
| Control 2 | 10X Chromium v2 | 2,000 | 88 | <sup>57</sup> |
| Control | Drop-Seq | 84 | 81 | <sup>58</sup> |
| Control | GemCode | 1,015 | 91 | <sup>52</sup> |
| Control | InDrops | 953 | 103 | <sup>59</sup> |

For each gene in each data set, we compared several expression models: a Gamma distribution, a point-Gamma distribution, a non-parametric unimodal distribution, and a fully non-parametric distribution (Figure 1a; Supplementary Note 2). Because these comparisons involve non-parametric families, obtaining *p*-values is not straightforward and perhaps inappropriate, since specifying any of these models as a “null” expression model is also questionable. Therefore, we instead compared the support for each model by comparing the likelihood of the data under each model. As a simple heuristic, we considered a likelihood ratio of 100 or more as strong evidence for one model over another.

**FIGURE 1.**
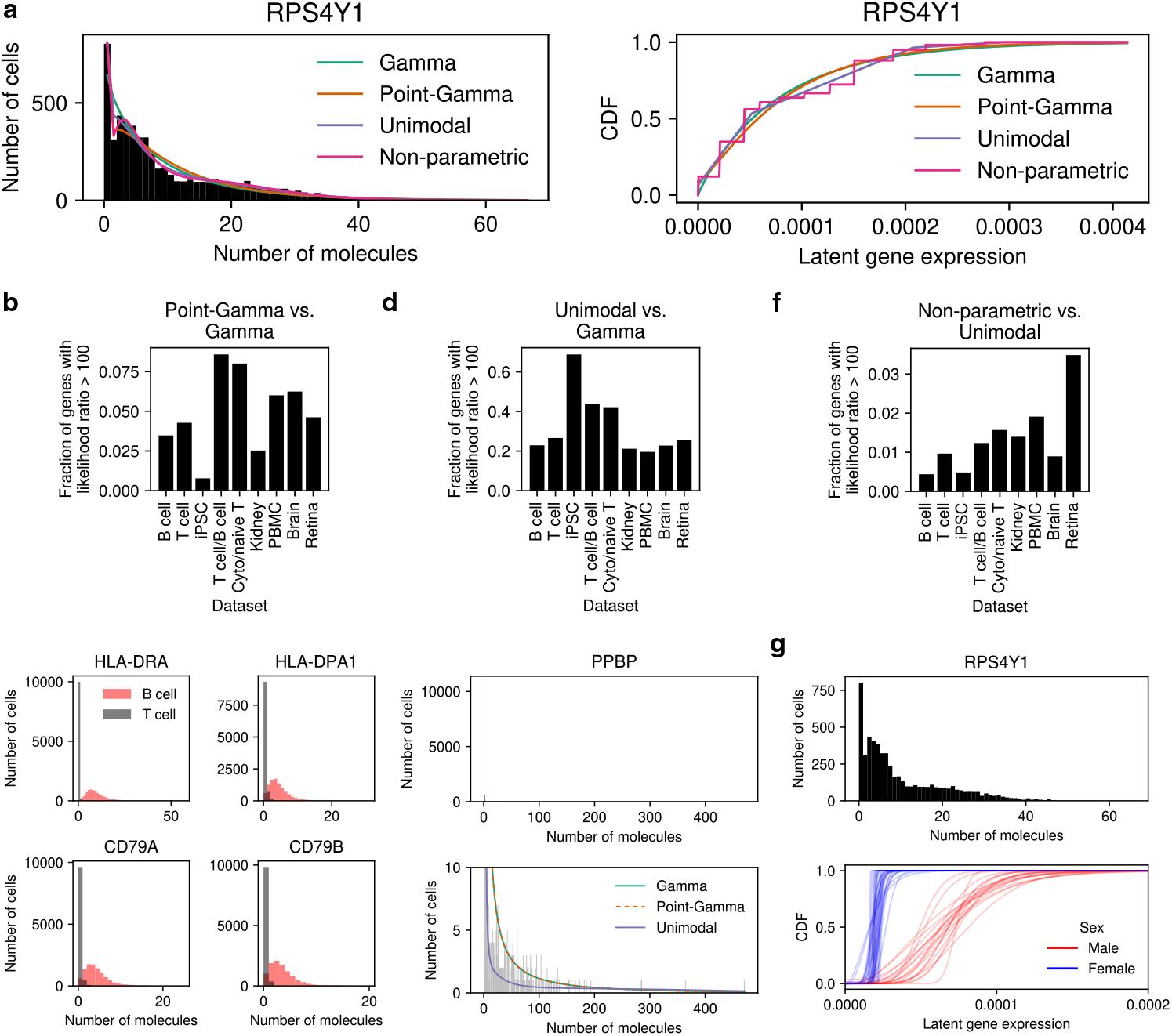
Comparing single-gene expression models on scRNA-seq data. (a) Example fits of different expression models, corresponding to different observation models, to observed data at a single gene. (b) Fraction of genes in biological data sets with strong evidence for a point-Gamma expression model over a Gamma model, and (c) examples of four genes showing strong evidence in an *in silico* mixture of T cells and B cells. (d) Fraction of genes with strong evidence in favor of a unimodal expression model over a Gamma model, and (e) an example of a gene showing strong evidence (*PPBP* in PBMCs). (f) Fraction of genes with strong evidence in favor of a fully non-parametric expression model over a unimodal non-parametric model, and (g) an example of a gene showing strong evidence (*RPS4Y1* in iPSCs).

We first assessed whether genes show evidence against a Gamma expression model due to excess zeros, by comparing a Gamma expression model with a point-Gamma model. In all biological data sets we examined, only a fraction of genes (0.6–9%) showed strong evidence in favor of the point-Gamma model (Figure 1b). The genes showing strong evidence in favor of the point-Gamma expression model included known marker genes in synthetic mixtures of sorted B cells and T cells (Figure 1c), providing a positive control that this approach can find such patterns.

Next, we assessed whether genes show evidence for other types of departures from a Gamma expression model, by comparing it to a non-parametric unimodal expression model. In this comparison, many more genes (20–69%) showed strong evidence in favor of the non-parametric unimodal expression model (Figure 1d). These results suggest that expression variation at many genes might not be captured even by a ZINB observation model. As an example, in PBMCs the gene *PPBP* exhibits not only many observed zeros, but also many small non-zero observations (for example, ones and twos) and a long tail of large observations (Figure 1e). Neither the Gamma nor point-Gamma distribution have sufficient flexibility to simultaneously describe both of these features, explaining the better fit of the non-parametric unimodal model.

Finally, we assessed whether the data show evidence of multimodal expression variation, by comparing a unimodal expression model against a fully non-parametric expression model. In this comparison, few genes (0.04–3%) showed strong evidence for the fully non-parametric expression model (Figure 1f), suggesting that multimodal expression variation may be rarer than previously suggested^27^. As a positive control, *RPS4Y1* is a Y chromosome-linked gene showing overwhelming evidence for the non-parametric expression model over the unimodal model (likelihood ratio > 10^65^), due to distinct distributions of gene expression in iPSCs derived from male and female donors (Figure 1g). One possible reason that cells from female donors are estimated to have non-zero expression of *RPS4Y1* is that the relevant part of the coding sequence is identical between *RPS4Y1* and its homolog *RPS4X*, and some reads erroneously mapped.

We emphasize that lack of multimodal expression variation does not imply lack of heterogeneity. To illustrate, *SKP1* is a gene at which expression in iPSCs is linked to the genotype of a nearby SNP, meaning both the mean and mode of expression levels vary across donors depending on genotype (Supplementary Figure 1). However, when the data are pooled across all samples, they show only modest evidence for multimodal expression variation (likelihood ratio 3.1 for the fully non-parametric expression model over the unimodal model). This is partly due to the substantial heterogeneity within each genotype class, which bridges the heterogeneity between genotype classes.

In summary, although some genes show departures from a simple Gamma model of expression variation, in most cases the data are consistent with unimodal expression variation, and relatively few genes show strong evidence for zero-inflated (point-Gamma) or multimodal expression variation.

#### Alternative measurement models

Our analysis above assumed a Poisson measurement model because it is both simple and has strong theoretical foundations. Here, we discuss the empirical support for the Poisson measurement model, as well as alternatives.

Our results on biological data sets above provide substantial evidence that the scRNA-seq measurement process is not zero-inflated. If the measurement process involved a distinct zero-generating component, then the observed data should have shown many genes supporting a point-Gamma expression model over a Gamma expression model; however, we did not find this. Indeed, only a fraction of genes (2–16%) show even suggestive evidence supporting a point-Gamma expression model (likelihood ratio > 10; Supplementary Figure 2). Therefore, we conclude that current data do not support the use of zero-inflated measurement models.

However, our results do not imply that the measurement model is necessarily Poisson. There are plausible reasons the measurement process could be overdispersed relative to Poisson; however, empirically assessing the extent of measurement overdispersion is difficult. In principle, one could analyze control data containing a known number of molecules of various synthetic genes. However, detecting subtle overdispersion would require very precise control of the number of molecules of each gene, which is difficult^4^. Without additional assumptions, it is not possible to say what extent overdispersion observed in control data is due to uncontrolled expression variation in the control genes versus measurement overdispersion.

To make progress on this question, we made an additional assumption that the measurement overdispersion is equal across genes. Under this assumption, it is possible to estimate the measurement dispersion by fitting an NB observation model with separate dispersion parameters to reflect measurement dispersion and uncontrolled expression variation (Supplementary Note 5).

We used this approach to bound the measurement dispersion in five control data sets (Table 3) that have been previously pre-processed and analyzed^4,16^. In all five data sets we found that the profile log likelihood for measurement dispersion dropped off quickly beyond some point, bounding the level that is consistent with the data (Supplementary Figure 3). Although the bounds vary among protocols, they suggest that measurement dispersion is no larger than 5 × 10^−3^ (likelihood ratio < 0.1 against the MLE). This level of measurement dispersion is small: for example, for a gene with mean 10 observed molecules, it increases the variance of the observations by 5% relative to a Poisson (and for genes with smaller mean, the increase in variance is smaller). Although this analysis does not rule out that measurement overdispersion could vary among genes, and could be large for some genes, we did not find empirical evidence for this.

### Multi-gene models

In multi-gene models, a common goal is to estimate the underlying low rank structure **LF’**, which could describe differences in the cell type/state of different samples, for example. Assessing the effectiveness of different methods for providing such biological insights is important, but also difficult to do objectively. For example, comparing clustering performance^28–30^ requires gold-standard labeled data which are arguably their ability to accurately estimate the underlying expression matrix, **Λ** = **LF’**. Surprisingly, we found that expression models making linear versus non-linear assumptions had largely similar accuracy on this problem (Supplementary Figure 4, Supplementary Note 3). The results suggest that practical issues (such as convergence behavior and computational cost), or subjective measures (such as ease of interpretation), may determine which methods are most useful in practice. Assessing expression models on these other metrics will be an important direction for future work.

## Conclusion

Here, we described how models for observed scRNA-seq counts can be helpfully separated into two parts: measurement models that describe variation introduced by the measurement process, and expression models that describe variation in true expression levels. We argued that a simple Poisson model is a reasonable starting point for the measurement model, and that many existing methods can be interpreted as combining a Poisson measurement model with different expression models. We explained how these simple ideas help clarify confusion about the source and interpretation of zeros in scRNA-seq data, and give rigorous procedures to interrogate variation in gene expression among cells.

How should one use these ideas in scRNA-seq analysis? We emphasize that clearly distinguishing between measurement, expression, and observation models can help reduce confusion and misinterpretation. In particular, both methods developers and data analysts should make explicit the assumptions made about measurement error and about expression variation when analyzing scRNA-seq data. One important area for future work will be developing fast and accurate diagnostics to assess whether these assumptions are violated by observed data, and whether analysis results are sensitive to these assumptions.

Interestingly, our empirical results suggest that simple expression models should suffice for common analysis tasks such as differential expression, dimension reduction, and clustering. Although we found that many genes showed support for a non-parametric unimodal expression model over the Gamma model, the Gamma model is considerably easier to fit and therefore may be preferred. Previous results have suggested that the impact of under-estimating expression variation on estimates of mean gene expression could be minimal^23^. Nonetheless, care may be necessary when dealing with long-tailed expression distributions, such as those exhibited by *PPBP* (Figure 1e).

Our empirical analyses of measurement error have two notable limitations. First, samples in spike-in experiments do not undergo the entire experimental protocol that biological samples do (dissociation, lysis, etc.), limiting their use to assess appropriate measurement models. This limitation is also shared by much prior work in this area^16,17,31–33^. Second, our approach to bound measurement dispersion made the strong assumption that measurement dispersion was equal across cells and genes. Therefore, the results may understate the potential for higher overdispersion at some genes. Despite these limitations, our empirical results clearly indicate that the use of zero-inflated measurement models is not supported.

There are commonly used methods that our framework does not encompass: for example, methods that first transform the count data (e.g. *y_ij_* = log(*x_ij_/x_i+_* + ϵ)) and then apply Gaussian methods such as principal components analysis^34^, factor analysis^35,^ or latent variable models^36,37^ to the transformed data **Y**. However, even for these methods, the key idea that observations reflect both expression variation and measurement error may still be useful to keep in mind. One potential way to formalize this is via Taylor series approximations^38,39^, for example E [*y_ij_* ≈ log (*λ_ij_* + ϵ), which suggest that Gaussian low rank models of the form E[**Y**] = **LF’** can be interpreted as assuming that [log (*λ_ij_* + ϵ)] is low rank. A key problem moving forward will be to make such connections rigorous and assess their accuracy.

## Acknowledgements

We thank members of the Stephens and Gilad labs for helpful comments. This work was supported by NIH grant HG002585 to M.S. and a Gut Cell Atlas grant from The Leona M. and Harry B. Helmsley Charitable Trust to M.S.

## Author Contributions

A.S. and M.S. developed the theory. A.S. performed the analysis. A.S. and M.S. wrote the paper.

## Box 1

### Assumptions and limitations of the Poisson measurement model

The Poisson measurement model (1) is based on the following assumptions (Supplementary Note 1): (i) in each cell, each molecule is equally likely to be observed, (ii) each molecule is observed independent of whether or not each other molecule is observed, and (iii) only a small proportion of all molecules present are observed. Assumptions (ii) and (iii) are plausible because the measurement process operates at the molecular level, and only 10–20% of molecules present are estimated to be observed in typical scRNA-seq experiments^7,60^. However, assumption (i) may plausibly be violated. There are many reasons a given mRNA molecule may fail to be observed: it can be lost to diffusion during sample collection and preparation, damaged by cell dissociation or lysis, or fail to be amplified or sequenced, for example. Different mRNA molecules will have different chances of surviving these processes, due to differences in RNA stability, location in the cell (e.g., nucleus vs cytoplasm), or sequence content, for example. Such factors could make the observed molecules a biased sample of all molecules.

Biased sampling of molecules can be incorporated into the Poisson measurement model by including bias terms (Supplementary Note 1). If the biases are systematic and associated with specific technical covariates (for example, batch) then one could estimate their effects within the Poisson model. However, some biases may vary from cell to cell in unknown ways (e.g., due to differences in the conditions under which each cell is processed). Such random biases effectively add additional noise to the measurement process, and could be dealt with by replacing the Poisson measurement model (1) with a negative binomial (NB) measurement model that allows overdispersion (additional variance) compared with Poisson. However, the NB measurement model raises additional difficulties, not least the question of how much overdispersion to allow for. Our perspective, for which we present empirical evidence, is that for many datasets the measurement overdispersion compared with Poisson will be small, especially compared with the variation in actual expression levels among cells, and so the Poisson measurement model (1) will often suffice.

Interestingly, as technologies improve to measure more molecules per cell – potentially violating assumption (iii) – the variance of the measurement process will be reduced relative to a Poisson (Supplementary Note 1). This could conceivably counteract, or even overshadow, some of the overdispersion mentioned above.

## Box 2

### Inference in the ZINB observation model

In our framework, the *Zero-inflated Negative Binomial* (ZINB) model for observations *x_1j_*, … , *x_nj_* is written

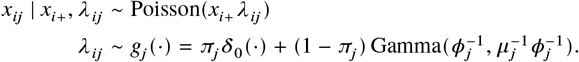

where *g_j_* is a *point-Gamma distribution*, which is a mixture of a point mass on zero (denoted δ_0_) and a Gamma distribution (parametrized by shape and rate). Here, we consider two analysis tasks: (1) estimating *g_j_* (i.e., estimating *π_j_, μ_j_, ϕ_j_*), and (2) estimating *λ_ij_*. Task (1) can be accomplished by maximizing the marginal likelihood, which has an analytic form but requires numerical optimization. From an estimate 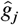, one can already estimate many useful quantities such as the mean and variance of gene expression^53^

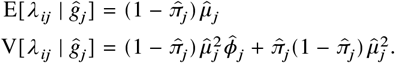

Task (2) can be accomplished by estimating the conditional distribution of *λ_ij_* given the observed data and 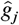

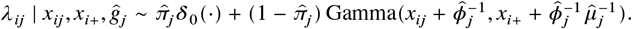

If one interprets *g_ij_* as a prior, then this procedure is empirical Bayes, and the mean of the conditional distribution above is the posterior mean estimate of *λ_ij_* given the observed data

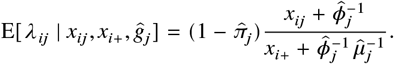

## Data Availability

Sorted immune cell and PBMC data were downloaded from https://10xgenomics.com/data. iPSC data were downloaded from Gene Expression Omnibus, accession number GSE118723. Brain data were downloaded from the GTEx portal https://www.gtexportal.org/home/datasets. Kidney and retina data were downloaded from the Human Cell Atlas Data Portal https://data.humancellatlas.org/. Control data were downloaded from https://figshare.com/projects/Zero_inflation_in_negative_control_data/61292. Analysis results generated in this study are available at https://zenodo.org/record/4543923 and analysis notebooks are available at https://aksarkar.github.io/singlecell-modes/.

## Code Availability

All code used to perform the analysis is available at https://zenodo.org/record/4543921 and https://zenodo.org/record/4543923.

## References

1. Fuller, W. A. Measurement Error Models (John Wiley & Sons, Inc., USA, 1986).

2. Marioni, J. C., Mason, C. E., Mane, S. M., Stephens, M. & Gilad, Y. RNA-seq: An assessment of technical reproducibility and comparison with gene expression arrays. Genome Res 18, 1509–1517 (2008).

3. Pachter, L. Models for transcript quantification from RNA-Seq. arXiv e-prints (2011).

4. Wang, J. et al. Gene expression distribution deconvolution in single-cell RNA sequencing. Proc Natl Acad Sci USA (2018).

5. Kharchenko, P. V., Silberstein, L. & Scadden, D. T. Bayesian approach to single-cell differential expression analysis. Nat Methods 11, 740 (2014).

6. Stegle, O., Teichmann, S. A. & Marioni, J. C. Computational and analytical challenges in single-cell transcriptomics. Nat Rev Genet 16. Review Article, 133 (2015).

7. Haque, A., Engel, J., Teichmann, S. A. & Lönnberg, T. A practical guide to single-cell RNA-sequencing for biomedical research and clinical applications. Genome Med 9, 75 (2017).

8. Zhu, L., Lei, J., Devlin, B. & Roeder, K. A unified statistical framework for single cell and bulk RNA sequencing data. Ann Appl Stat 12, 609–632 (2018).

9. Qiu, P. Embracing the dropouts in single-cell RNA-seq analysis. Nat Commun 11, 1169 (2020).

10. Fujimura, F. K., Northrup, H., Beaudet, A. L. & O’Brien, W. E. Genotyping Errors with the Polymerase Chain Reaction. N Engl J Med 322, 61–61 (1990).

11. Whale, A. S., Cowen, S., Foy, C. A. & Huggett, J. F. Methods for Applying Accurate Digital PCR Analysis on Low Copy DNA Samples. PLoS ONE 8, 1–10 (2013).

12. Hicks, S. C., Townes, F. W., Teng, M. & Irizarry, R. A. Missing data and technical variability in single-cell RNA-sequencing experiments. Biostatistics (2017).

13. Li, W. V. & Li, J. J. An accurate and robust imputation method scImpute for single-cell RNA-seq data. Nat Commun 9, 997 (2018).

14. Chen, M. & Zhou, X. VIPER: variability-preserving imputation for accurate gene expression recovery in single-cell RNA sequencing studies. Genome Biol 19, 196 (2018).

15. Talwar, D., Mongia, A., Sengupta, D. & Majumdar, A. AutoImpute: Autoencoder based imputation of single-cell RNA-seq data. Sci Rep 8, 16329 (2018).

16. Svensson, V. Droplet scRNA-seq is not zero-inflated. Nat Biotech (2020).

17. Islam, S. et al. Quantitative single-cell RNA-seq with unique molecular identifiers. Nat Methods 11, 163 (2013).

18. Townes, F. W., Hicks, S. C., Aryee, M. J. & Irizarry, R. A. Feature selection and dimension reduction for single-cell RNA-Seq based on a multinomial model. Genome Biol 20, 295 (2019).

19. Huang, M. et al. SAVER: gene expression recovery for single-cell RNA sequencing. Nat Methods 15, 539–542 (2018).

20. Eraslan, G., Simon, L. M., Mircea, M., Mueller, N. S. & Theis, F. J. Single-cell RNA-seq denoising using a deep count autoencoder. Nat Commun 10, 390 (2019).

21. Tang, W. et al. bayNorm: Bayesian gene expression recovery, imputation and normalization for single-cell RNA-sequencing data. Bioinformatics 36, 1174–1181 (2019).

22. Hilbe, J. M. Modeling Count Data (Cambridge University Press, 2014).

23. Lu, M. Generalized Adaptive Shrinkage Methods and Applications in Genomics Studies PhD thesis (University of Chicago, 2018).

24. Raj, A. & van Oudenaarden, A. Nature, Nurture, or Chance: Stochastic Gene Expression and Its Consequences. Cell 135, 216–226 (2008).

25. Shalek, A. K. et al. Single-cell transcriptomics reveals bimodality in expression and splicing in immune cells. Nature 498, 236 (2013).

26. Shalek, A. K. et al. Single-cell RNA-seq reveals dynamic paracrine control of cellular variation. Nature 510, 363 (2014).

27. Bacher, R. & Kendziorski, C. Design and computational analysis of single-cell RNA-sequencing experiments. Genome Biol 17, 63 (2016).

28. Lopez, R., Regier, J., Cole, M. B., Jordan, M. I. & Yosef, N. Deep generative modeling for single-cell transcriptomics. Nat Methods 15, 1053–1058 (2018).

29. Hu, Q. & Greene, C. S. Parameter tuning is a key part of dimensionality reduction via deep variational autoencoders for single cell RNA transcriptomics. Pac Symp Biocomput 24, 362–373 (2019).

30. Sun, S., Zhu, J., Ma, Y. & Zhou, X. Accuracy, robustness and scalability of dimensionality reduction methods for single-cell RNA-seq analysis. Genome Biol 20, 269 (2019).

31. Brennecke, P. et al. Accounting for technical noise in single-cell RNA-seq experiments. Nat Methods 10, 1093 (2013).

32. Kim, J. K., Kolodziejczyk, A. A., Ilicic, T., Teichmann, S. A. & Marioni, J. C. Characterizing noise structure in single-cell RNA-seq distinguishes genuine from technical stochastic allelic expression. Nature Commun 6, 8687 (2015).

33. Wang, W. & Stephens, M. Empirical Bayes Matrix Factorization. arXiv e-prints (2018).

34. Tipping, M. E. & Bishop, C. M. Probabilistic Principal Component Analysis. Journal of the Royal Statistical Society: Series B (Statistical Methodology) 61, 611–622.

35. Pierson, E. & Yau, C. ZIFA: Dimensionality reduction for zero-inflated single-cell gene expression analysis. Genome Biol 16, 241 (2015).

36. Buettner, F. et al. Computational analysis of cell-to-cell heterogeneity in single-cell RNA-sequencing data reveals hidden subpopulations of cells. Nat Biotech 33, 155–160 (2015).

37. Verma, A. & Engelhardt, B. E. A robust nonlinear low-dimensional manifold for single cell RNA-seq data. BMC Bioinformatics 21, 324 (2020).

38. Law, C. W., Chen, Y., Shi, W. & Smyth, G. K. voom: precision weights unlock linear model analysis tools for RNA-seq read counts. Genome Biol 15, R29 (2014).

39. Lun, A. Overcoming systematic errors caused by log-transformation of normalized single-cell RNA sequencing data. bioRxiv (2018).

40. Venables, W. N. & Ripley, B. D. Modern Applied Statistics with S Fourth. ISBN 0-387-95457-0 (Springer, New York, 2002).

41. Robinson, M. D., McCarthy, D. J. & Smyth, G. K. edgeR: a Bioconductor package for differential expression analysis of digital gene expression data. Bioinformatics 26, 139–140 (2009).

42. Love, M. I., Huber, W. & Anders, S. Moderated estimation of fold change and dispersion for RNA-seq data with DESeq2. Genome Biol 15, 550 (2014).

43. Vallejos, C. A., Marioni, J. C. & Richardson, S. BASiCS: Bayesian Analysis of Single-Cell Sequencing Data. PLoS Comp Biol 11, 1–18 (2015).

44. Zeileis, A., Kleiber, C. & Jackman, S. Regression Models for Count Data in R. Journal of Statistical Software 27 (2008).

45. Stephens, M. False discovery rates: a new deal. Biostatistics 18, 275–294 (2017).

46. Kiefer, J. & Wolfowitz, J. Consistency of the Maximum Likelihood Estimator in the Presence of Infinitely Many Incidental Parameters. Ann Math Statist 27, 887–906 (1956).

47. Lee, D. D. & Seung, H. S. Algorithms for Non-negative Matrix Factorization in Advances in Neural Information Processing Systems 13, Papers from Neural Information Processing Systems (NIPS) 2000, Denver, CO, USA (eds Leen, T. K., Dietterich, T. G. & Tresp, V.) (MIT Press, 2000), 556–562.

48. Levitin, H. M. et al. De novo gene signature identification from single-cell RNA-seq with hierarchical Poisson factorization. Mol Syst Biol 15 (2019).

49. Gouvert, O., Oberlin, T. & Févotte, C. Negative Binomial Matrix Factorization for Recommender Systems. arXiv e-prints (2018).

50. Sun, S., Chen, Y., Liu, Y. & Shang, X. A fast and efficient count-based matrix factorization method for detecting cell types from single-cell RNAseq data. BMC Syst Biol 13, 28 (2019).

51. Risso, D., Perraudeau, F., Gribkova, S., Dudoit, S. & Vert, J.-P. A general and flexible method for signal extraction from single-cell RNA-seq data. Nat Commun 9, 284 (2018).

52. Zheng, G. X. Y. et al. Massively parallel digital transcriptional profiling of single cells. Nat Commun 8, 14049 (2017).

53. Sarkar, A. K. et al. Discovery and characterization of variance QTLs in human induced pluripotent stem cells. PLoS Genetics 15, 1–16 (2019).

54. Habib, N. et al. Massively parallel single-nucleus RNA-seq with DroNc-seq. Nature Methods 14, 955–958 (2017).

55. Stewart, B. J. et al. Spatiotemporal immune zonation of the human kidney. Science 365, 1461–1466 (2019).

56. Lukowski, S. W. et al. A single-cell transcriptome atlas of the adult human retina. The EMBO Journal 38, e100811 (2019).

57. Svensson, V. et al. Power analysis of single-cell RNA-sequencing experiments. Nature Methods 14, 381–387 (2017).

58. Macosko, E. Z. et al. Highly Parallel Genome-wide Expression Profiling of Individual Cells Using Nanoliter Droplets. Cell 161, 1202–1214 (2015).

59. Klein, A. M. et al. Droplet Barcoding for Single-Cell Transcriptomics Applied to Embryonic Stem Cells. Cell 161, 1187–1201 (2015).

60. Islam, S. et al. Characterization of the single-cell transcriptional landscape by highly multiplex RNA-seq. Genome Res 21, 1160–1167 (2011).

