## Supplementary Information for "Separating measurement and expression models clarifies confusion in single-cell RNA sequencing analysis"

### SUPPLEMENTARY INFORMATION FOR "SEPARATING MEASUREMENT AND EXPRESSION MODELS CLARIFIES CONFUSION IN SINGLE CELL RNA-SEQ ANALYSIS"

ABHISHEK SARKAR AND MATTHEW STEPHENS

#### Contents

1. Poisson model for measurement error 2
2. Inference in single-gene models 4
3. Inference in multi-gene models 6
4. Identifiability of measurement and expression models 9
5. Negative binomial model for measurement error 10
6. Supplementary Methods 11
7. Supplementary Figures 13

References 16

#### 1. Poisson model for measurement error

##### 1.1. **Derivation of basic Poisson measurement model.** 
Let $m_{ij}$ denote the number of mRNA molecules present in cell $i$ from gene $j$ ($i = 1, \dots, n; j = 1, \dots, p$), and let $m_{i+} \triangleq \sum_j m_{ij}$. The simplest model for measurement error is to assume that the measured molecules in each cell are a simple random sample of all available molecules (sampled without replacement through the use of UMIs). Under this assumption, the observed molecule counts for each gene in sample $i$, $x_{i1}, \dots, x_{ip}$, follow a Multivariate Hypergeometric distribution. Unfortunately, the Multivariate Hypergeometric distribution is difficult to work with. However, if the number of molecules sampled $x_{i+} \triangleq \sum_j x_{ij}$ is small compared with the total number of molecules in the cell ($x_{i+} \ll m_{i+}$), then it is reasonable and convenient to approximate the Multivariate Hypergeometric distribution with the simpler Multinomial distribution (i.e., assuming sampling with replacement):

$$x_{i1}, \dots, x_{ip} \mid x_{i+}, \lambda_{i1}, \dots, \lambda_{ip} \sim \text{Multinomial}(x_{i+}; \lambda_{i1}, \dots, \lambda_{ip}), \tag{1}$$

where $\lambda_{ij} \triangleq m_{ij}/m_{i+}$ is the relative expression of gene $j$ in sample $i$. Furthermore, it is often still more convenient to transform (1) into a Poisson model[1,2]:

$$x_{ij} \mid x_{i+}, \lambda_{ij} \sim \text{Poisson}(x_{i+}\lambda_{ij}), \qquad j = 1, \dots, p. \tag{2}$$

It is straightforward to show that the models (1) and (2) are equivalent, meaning they produce the same (log) likelihood for $\lambda_{i\cdot}$ up to a constant, which does not affect either maximum likelihood or Bayesian inference. Let $l_{\text{pois}}$ and $l_{\text{mult}}$ denote the log-likelihoods for $\lambda_{i\cdot}$ under (2) and (1), respectively. Then,

$$l_{\text{pois}}(\lambda_{i\cdot}) \triangleq \ln p(x_{i1}, \dots, x_{ip} \mid x_{i+}, \lambda_{i\cdot}) \tag{3}$$

$$= \sum_j \left[ x_{ij} \ln(x_{i+}\lambda_{ij}) - x_{i+}\lambda_{ij} \right] + \text{const} \tag{4}$$

$$= \sum_j x_{ij} \ln \lambda_{ij} + \text{const} \qquad (\text{because } \textstyle\sum_j \lambda_{ij} = 1) \tag{5}$$

$$= l_{\text{mult}}(\lambda_{i\cdot}) + \text{const}. \tag{6}$$

##### 1.2. **Incorporating non-random sampling.** 
The derivation above assumed random sampling of molecules, with all molecules equally likely to be sampled. However, as outlined in the main text, a number of technical effects might violate this assumption, either by making some molecules more likely to be sampled than others, or by making observing one molecule dependent on observing other molecules.

Perhaps the simplest type of non-random sampling is biased sampling, where molecules are sampled independently of one another, but some molecules are more likely to be sampled than others. Biased sampling can be incorporated into the Poisson measurement model (2) by adding bias terms $b_{ij}$ that multiply the relative expression levels $\lambda_{ij}$. To make this idea more precise, suppose that the probability of successfully sampling a molecule from gene $j$ in cell $i$ is proportional to $b_{ij}$, and suppose the measurement process generates each measured molecule by repeatedly drawing from the pool of molecules present until the first success. Then, for each molecule in cell $i$

$$\Pr(\text{measured molecule from gene } j \mid \text{measurement succeeded}) = \frac{b_{ij} m_{ij}}{\sum_{it} b_{it} m_{it}}. \tag{7}$$

Without loss of generality, scale $b_{i1}, \dots, b_{ip}$ such that $\sum_j b_{ij} m_{ij} = m_{i+}$. Then,

$$x_{i1}, \dots, x_{ip} \mid x_{i+}, \lambda_{i1}, \dots, \lambda_{ip}, b_{i1}, \dots, b_{ip} \sim \text{Multinomial}(x_{i+}; b_{i1}\lambda_{i1}, \dots, b_{ip}\lambda_{ip}), \tag{8}$$

implying

$$x_{ij} \mid x_{i+}, \lambda_{ij}, b_{ij} \sim \text{Poisson}(x_{i+} b_{ij} \lambda_{ij}), \qquad j = 1, \dots p. \tag{9}$$

A similar argument was previously made in modeling the impact of gene length on the measurement model for bulk RNA-seq read counts[2]. In some cases, it may be possible to estimate the values of these bias terms from observed data, for example when the biases arise from observed technical covariates for each cell[3,4]. However, it will be impractical to identify and treat all possible sources of bias in this way.

What happens if one ignores biased sampling, and uses the model (2) as is? In this case, the measurement model describes not the true expression levels $\lambda_{ij}$ but instead $\tilde{\lambda}_{ij} = b_{ij}\lambda_{ij}$. If the bias terms $b_{ij}$ are the same for every cell, then the ratio $\tilde{\lambda}_{ij}/\tilde{\lambda}_{i'j}$ for two different cells $i$ and $i'$ will equal the ratio of the true expression levels $\lambda_{ij}/\lambda_{i'j}$, because the bias terms cancel. However, if biases vary randomly from cell to cell, then they will lead to overdispersed measurement error relative to Poisson.

A more complex type of non-random sampling is dependent sampling of molecules. For example, which molecules remain undamaged and successfully adhere to barcoded beads (in droplet-based platforms) after cell lysis may depend on their initial spatial location within the cell. Dependent sampling of molecules (in addition to unaccounted-for biases described above) will lead to the measurements having larger variance than a Poisson distribution – that is, being overdispersed relative to Poisson. In principle, measurement overdispersion could be incorporated by replacing the Poisson measurement model with an NB measurement model. However, doing so introduces complications, not least the difficulty of knowing how much overdispersion to attribute to the measurement process. Precisely characterizing the overdispersion of the measurement process from empirical data is not straightforward, and requires additional assumptions (see Supplementary Note 5 for details). What happens if one ignores overdispersed measurement error, and analyzes data using the Poisson measurement model? In brief, such analyses may tend to overstate the variability of expression levels among cells, because they will wrongly ascribe some of the additional variance from the measurement process to expression variation.

1.3. **Sampling a large fraction of molecules.** The derivations above used a Multinomial (Poisson) model to approximate the Multivariate Hypergeometric distribution, which relies on the assumption $x_{i+} \ll m_{i+}$. We now briefly consider what could happen if this assumption is violated; i.e., if one actually observed a substantial fraction of all available molecules.

Consider gene $j$, and partition the molecules into those that map to gene $j$ and those that do not. Then, the Multivariate Hypergeometric distribution reduces to the Hypergeometric distribution, and the Multinomial distribution reduces to the Binomial distribution. Suppose one sequences a fraction $p$ of the molecules, and consider the following candidate models:

$$x_{ij} \sim \text{Poisson}(m_{ij} p) \triangleq P_1 \tag{10}$$

$$x_{ij} \sim \text{Binomial}(m_{i+} p, m_{ij}/m_{i+}) \triangleq P_2 \tag{11}$$

$$x_{ij} \sim \text{Hypergeometric}(m_{i+} p, m_{i+}, m_{ij}) \triangleq P_3. \tag{12}$$

These models all have the same expectation, $\mathrm{E}[x_{ij}] = m_{ij}p$, but unequal variances:

$$\mathrm{V}_{P_1}[x_{ij}] = m_{ij}p \tag{13}$$

$$\mathrm{V}_{P_2}[x_{ij}] = m_{ij}p\frac{m_{i+} - m_{ij}}{m_{i+}} \tag{14}$$

$$\mathrm{V}_{P_3}[x_{ij}] = m_{ij}p\frac{m_{i+} - m_{ij}}{m_{i+}}\frac{m_{i+}}{m_{i+} - 1}(1 - p). \tag{15}$$

It is reasonable to assume that $m_{i+}$ is large ($m_{i+} >> m_{ij}$ and $m_{i+} >> 1$), in which case both $(m_{i+} - m_{ij})/m_{i+}$ and $m_{i+}/(m_{i+} - 1)$ are close to one. Therefore, the difference in variance between the Poisson and Hypergeometric measurement model is primarily due to the factor $1 - p$. By definition, $p \leq 1$; therefore, the Poisson measurement model has larger variance than the Hypergeometric measurement model and will tend to overstate the amount of measurement error as $p$ increases. If technologies develop to sample very large fractions of molecules ($p \approx 1$) then it is possible that statistical methods could benefit from taking the resulting reduced measurement variance into account.

Note that the effects of sampling a large proportion of molecules work in opposition to the factors considered in the previous section: whereas the biases in the previous section tend to increase measurement dispersion relative to Poisson, the sampling of large proportions of molecules will tend to decrease measurement dispersion relative to Poisson. The balance between these factors is difficult to assess, which we view as an argument for keeping models simple – that is, starting with a Poisson measurement model rather than rushing to incorporate overdispersion into the measurement process.

#### 2. Inference in single-gene models

To analyze observed data $x_{1j}, \dots, x_{nj}$ at a single gene $j$, one combines the Poisson measurement model with an expression model $g_j$

$$\begin{aligned} x_{ij} \mid x_{i+}, \lambda_{ij} &\sim \mathrm{Poisson}(x_{i+}\lambda_{ij}) \\ \lambda_{ij} &\sim g_j(\cdot) \in \mathcal{G}, \end{aligned} \tag{16}$$

where $\mathcal{G}$ is some family of distributions, and estimates $g_j$ from the data. This is achieved by maximizing the likelihood

$$L(g_j; x_{1j}, \dots, x_{nj}, x_{1+}, \dots, x_{n+}) \triangleq \prod_{i=1}^{n} \int_0^\infty \mathrm{Poisson}(x_{ij}; x_{i+}\lambda_{ij})\, dg_j(\lambda_{ij}) \tag{17}$$

$$\hat{g}_j = \arg\max_{g_j \in \mathcal{G}} L(g_j; x_{1j}, \dots, x_{nj}, x_{1+}, \dots, x_{n+}). \tag{18}$$

Readers who are more familiar with likelihoods for parametric distributions should not be put off by our notation $L(g_j)$ here, which is simply the usual definition of likelihood, but with the "parameter" being a distribution. If $\mathcal{G}$ is a parametric family, parameterized by $\theta \in \Theta$ say, then maximizing $L(g_j)$ over $g_j \in \mathcal{G}$ is simply the same as maximizing $L(\theta)$ over $\theta \in \Theta$. We use the more general notation to allow that $\mathcal{G}$ could be a non-parametric family.

2.1. **Methods for selected expression models.** Different assumptions on $\mathcal{G}$ trade off ease of implementation and computational cost for flexibility and modeling power. If one makes strong parametric assumptions about $\mathcal{G}$, then the estimation procedures are easy. In contrast, more flexible models require more sophisticated numerical optimization algorithms to estimate from observed data.

2.1.1. *Point mass expression model.* If one assumes that the true gene expression levels $\lambda_{ij} = \mu_j$, i.e., that there is no variation in expression levels, then the MLE

$$\hat{\mu}_j = \frac{\sum_i x_{ij}}{\sum_i x_{i+}}. \tag{19}$$

2.1.2. *(Point-)Gamma expression model.* If one assumes $g_j$ is a Gamma distribution, then the integrals in (17) are analytic, and yield Negative Binomial distributions. (We highlight a subtle point: since the measurement model for $x_{ij}$ depends on $x_{i+}$, each observation comes from its own specific observation model $p(x_{ij} \mid x_{i+}, g_j)$.) Similarly, if one assumes $g_j$ is a mixture of a point mass on zero and a Gamma distribution, then the integrals yield Zero-inflated Negative Binomial distributions. In the simplest cases, these observation models can be written as generalized linear models, and can be readily fit using standard packages[5,6]. They can also be fit using EM algorithms[7,8]. Additionally, many published methods for both bulk and single cell RNA-seq data have adopted the NB observation model, although they do not necessarily directly perform downstream inferences on the corresponding Gamma expression model.

2.1.3. *Unimodal expression model.* If one assumes $g_j$ is a unimodal (non-parametric) distribution on non-negative reals, then inference can be performed using *generalized adaptive shrinkage*[9,10]. In this approach, $g_j$ is represented as a large, but finite, mixture of uniform distributions

$$g_j(\cdot) = \sum_{k=1}^{K_j} w_{jk} \,\mathrm{Uniform}(\cdot; \lambda_{0j}, a_{jk}), \tag{20}$$

where the number of endpoints $K_j$ and their locations $a_{jk}$ are fixed based on the range of the data. (Here $\mathrm{Uniform}(\cdot; \lambda_{0j}, a_{jk})$ denotes the density of the uniform distribution on $[a_{jk}, \lambda_{0j}]$ if $a_{jk} < \lambda_{0j}$, and on $[\lambda_{0j}, a_{jk}]$ if $a_{jk} > \lambda_{0j}$.) Assuming the mode $\lambda_{0j}$ is known, the likelihood (17) is a function only of the mixture weights $\mathbf{w}_j$, resulting in a convex optimization problem. To estimate $\lambda_{0j}$, Brent's method is used to optimize (17) with respect to $\lambda_{0j}$, at each candidate point optimizing over mixture weights $\mathbf{w}_j$. This approach is implemented in the R package *ashr*.

2.1.4. *Point-exponential family expression model.* If one assumes $g_j$ is a mixture of a point mass on zero and an exponential family, then inference can performed by numerically optimizing the marginal likelihood[3,11]. In this approach,

$$g_j(\cdot) = \pi_j \delta_0(\cdot) + (1 - \pi_j) \exp\left(\mathbf{Q}(\cdot)'\boldsymbol{\alpha} - \phi(\boldsymbol{\alpha})\right) \tag{21}$$

where $\mathbf{Q}(\cdot)$ is the degree 5 natural cubic spline basis, $\boldsymbol{\alpha}$ is the $5 \times 1$ natural parameter, and $\phi(\cdot)$ is the normalizer. In practice, the basis $\mathbf{Q}$ and the representation of the distribution $g_j$ are both discretized, and a regularization penalty is added to the likelihood (17). This approach is flexible enough to represent many unimodal distributions and some multi-modal distributions due to its use of (smooth) natural splines, and is implemented in the R package *DESCEND*.

2.1.5. *Fully non-parametric expression model.* The most general approach is to assume that $g_j$ is simply some (non-parametric) distribution on non-negative reals[12,13]. Any such distribution $g_j$ can be approximated by a mixture of uniforms on a dense grid

$$g_j(\cdot) = \sum_{k=1}^{K_j} w_{jk} \mathrm{Uniform}(\cdot; (k-1)a_j, ka_j), \tag{22}$$

where the number of components $K_j$ depends on the range of the data and the grid step size $a_j$ is fixed. (Alternatively, one could fix $K_j$, and choose $a_j$ depending on the range of the data.) Fitting this expression model reduces to the same convex optimization problem as (20), and is implemented in the R package *ashr*. In practice, we begin with a relatively large step size $a_j$, and iteratively discard grid segments with weight $w_{jk}$ close to zero and refine (split) the remaining grid segments, until the improvement in log likelihood becomes smaller than a threshold. This iterative refinement approach is implemented in the Python package *scmodes* (`https://www.github.com/aksarkar/scmodes`).

2.2. **Incorporating dependence between relative expression levels and size factors.** Under the combined model (16), relative gene expression levels $\lambda_{ij}$ are assumed to be independent of the total number of molecules observed $x_{i+}$. However, this assumption could be violated in real data, because e.g., the total number of molecules observed depends in part on the total number present. Therefore, the independence assumption could be violated if e.g., the total number of molecules present depends on cell type/state and the gene $j$ is differentially expressed in different cell types/states. In this case, the fitted observation model may not adequately describe the data, regardless of how flexible $\mathcal{G}$ is.

To relax the independence assumption, let $\tilde{\lambda}_{ij} \triangleq x_{i+}^{\alpha} \lambda_{ij}$, where $0 \leq \alpha \leq 1$, and consider the measurement and expression models

$$
\begin{aligned}
x_{ij} \mid x_{i+}, \tilde{\lambda}_{ij} &\sim \operatorname{Poisson}(x_{i+}^{1-\alpha} \tilde{\lambda}_{ij}) \\
\tilde{\lambda}_{ij} &\sim g_j(\cdot) \in \mathcal{G}.
\end{aligned} \tag{23}
$$

For fixed $\alpha$, the inference scheme presented above (including specific algorithms for specific choices of $\mathcal{G}$) can be applied to observations $x_{1j}, \ldots, x_{nj}$ and size factors $x_{1+}^{1-\alpha}, \ldots, x_{n+}^{1-\alpha}$ to estimate $g_j$. The resulting estimate $\hat{g}_j$ is a prior on $\tilde{\lambda}_{ij}$, and can be transformed into the prior on relative expression levels $p(\lambda_{ij})$, and in turn used for posterior inference.

Based on this fact, $\alpha$ can be estimated by maximizing the likelihood (16) using e.g., Brent's method. For specific choices of $\mathcal{G}$, more efficient methods are possible. For example, assuming $g_j$ is a Gamma distribution, $\alpha$ can be estimated by a negative binomial GLM with design matrix equal to the (log-transformed) vector $x_{1+}, \ldots, x_{n+}$ and an intercept $\mu_j$ (the mean of the Gamma expression model).

#### 3. Inference in multi-gene models

To analyze variation in the observed data $\mathbf{X}$ across both cells and genes simultaneously, one combines the Poisson measurement model with an expression model of the form

$$
\begin{aligned}
\lambda_{ij} &= \mu_{ij} u_{ij} \\
\mu_{ij} &= h^{-1}((\mathbf{L}\mathbf{F}')_{ij}) \\
u_{ij} &\sim p(u_{ij}),
\end{aligned} \tag{24}
$$

where $h$ is a *link function*, or transformation; the *loadings* matrix $\mathbf{L}$ is $n \times k$; the *factors* matrix $\mathbf{F}$ is $p \times k$ ($\mathbf{F}'$ indicates the transpose of the matrix $\mathbf{F}$); and $p(u_{ij})$ denotes an assumed distribution of stochastic deviations from the low rank structure. This model can equivalently be understood as assuming $\mu_{i\cdot} = f(l_{i\cdot})$, where $f$ is a function, parameterized by $\mathbf{F}$, to be learned from the data. The model (24) can be further generalized by e.g., assuming $f$ is parameterized by a neural network, or allowing $p(u_{ij})$ to vary with $i$ and/or $j$ (e.g., allowing gene-specific and/or sample-specific variance).

In principle, one could incorporate a link function in a single-gene expression model $h(\lambda_{ij}) \sim g_j(\cdot) \in \mathcal{G}$; however, doing so would be exactly equivalent to transforming the distribution $g_j$, and so would not provide any additional generality. In contrast, assuming $h$ is identity in (24) is fundamentally different from assuming $h = \log$: if the matrix $[\lambda_{ij}]$ is rank $k$, it is not necessarily true that the matrix $[\log \lambda_{ij}]$ is also rank $k$ (or vice versa).

3.1. **Methods for selected multi-gene expression models.** Marginalizing over $u_{ij}$ yields $\lambda_{ij} \sim g_{ij}(\cdot) \in \mathcal{G}$, connecting single-gene and multi-gene expression models in our framework. Therefore, fitting multi-gene expression models to data involves the same two steps as fitting single-gene expression models: (1) estimating $g$ by maximizing the marginal likelihood, and (2) estimating $\lambda_{ij}$ (or $\mu_{ij}$, in models with noise).

3.1.1. *Poisson Non-negative Matrix Factorization.* If one assumes that the entries of $\mathbf{L}$ and $\mathbf{F}$ are non-negative, and that $u_{ij} = 1$, then one obtains the Poisson NMF observation model[14,15]. The model can be fit by minimizing a generalized KL divergence by multiplicative updates[16], which is equivalent to maximizing the Poisson log likelihood by expectation maximization[17].

3.1.2. *GLM-PCA.* If one assumes $h = \ln$ and that $u_{ij} = 1$, then one obtains the GLM-PCA observation model[18]. The implementation of GLM-PCA also supports the case where $u_{ij} \sim \mathrm{Gamma}(\phi_j^{-1}, \phi_j^{-1})$, where the Gamma distribution is parameterized by shape and rate (i.e., with mean one and variance $\phi_j$). The model is fit by maximizing the likelihood using Fisher scoring (Newton-Raphson updates, using the expected information matrix in place of the Hessian), and is implemented in the R package *glmpca*.

3.1.3. *ZINB-WaVE.* If one assumes $u_{ij}$ follows a point-Gamma distribution

$$u_{ij} \mid \pi_{ij}, \phi_{ij} \sim \pi_{ij}\delta_0(\cdot) + (1 - \pi_{ij})\,\mathrm{Gamma}(\phi_{ij}^{-1}, \phi_{ij}^{-1}), \tag{25}$$

where the Gamma distribution is parameterized by shape and rate (i.e., with mean one and variance $\phi_{ij}$), and additionally assumes that

$$\ln \mu_{ij} = (\mathbf{L}\mathbf{F}'_\mu)_{ij} \tag{26}$$

$$\phi_{ij} = \phi_j \tag{27}$$

$$\operatorname{logit} \pi_{ij} = (\mathbf{L}\mathbf{F}'_\pi)_{ij}, \tag{28}$$

then one obtains the ZINB-WaVE observation model[19]. (For brevity, we have omitted fixed effects for observed sample- and gene-specific covariates.) We note that in this formulation, $p(u_{ij})$ depends on $l_{i\cdot}$, complicating our proposed interpretation of $u_{ij}$ as capturing expression noise. The model is fit by maximizing a penalized log likelihood using alternating optimization, and is implemented in the R package *zinbwave*. We note that the description here differs from the original presentation of the method, with consequences on downstream inference (see Supplementary Note 4 for details).

3.1.4. *DCA.* If one assumes that $u_{ij}$ follows a point-Gamma distribution (25), and that $f$ is a neural network mapping a $k$-dimensional representation $l_{i\cdot}$ to parameters of $g_{ij}$, then one obtains the DCA observation model[20]. Specifically, $f$ produces outputs

$$\mu_{i\cdot}(l_{i\cdot}) = \exp(\mathbf{W}_\mu \,\mathrm{ReLU}(\mathbf{W}_\mathbf{L} l'_{i\cdot})) \tag{29}$$

$$\phi_{i\cdot}(l_{i\cdot}) = \exp(\mathbf{W}_\phi \,\mathrm{ReLU}(\mathbf{W}_\mathbf{L} l'_{i\cdot})) \tag{30}$$

$$\pi_{i\cdot}(l_{i\cdot}) = \mathrm{sigmoid}(\mathbf{W}_\pi \,\mathrm{ReLU}(\mathbf{W}_\mathbf{L} l'_{i\cdot})), \tag{31}$$

where $\mathrm{sigmoid}(x) = 1/(1 + \exp(-x))$; $\mathrm{ReLU}(x) = \max(0, x)$; functions are applied element-wise; $\mathbf{W}_\mathbf{L}$ is an $m \times k$ matrix, where $k < m < p$; and $\mathbf{W}_\mu, \mathbf{W}_\phi, \mathbf{W}_\pi$ are $p \times m$

matrices. We note that in this formulation, $p(u_{ij})$ depends on $l_{i\cdot}$, complicating our proposed interpretation of $u_{ij}$ as capturing expression noise.

In practice, inference is performed by introducing an *inference network*[21–24], which is a neural network $\tilde{f}$ mapping $x_{i\cdot}$ to $l_{i\cdot}$, and minimizing the ZINB negative log likelihood by stochastic gradient descent. This algorithm is implemented in the Python package *dca*. (The software implementation of DCA also supports a Poisson observation model, $u_{ij} = 1$, and an NB observation model, $u_{ij} \sim \text{Gamma}(\phi_{ij}^{-1}, \phi_{ij}^{-1})$.) We note that the description here differs from the original presentation of the method, with consequences on downstream inference (see Supplementary Note 4 for details).

3.1.5. *scVI.* If one assumes that $u_{ij}$ follows a point-Gamma distribution (25), that $\phi_{ij} = \phi_j$ (in the simplest case), that $f$ is a neural network mapping a $k$-dimensional representation $l_{i\cdot}$ to parameters of $g_{ij}$ (termed a *decoder network*[22]), and that $l_{i\cdot} \sim \text{Normal}(\mathbf{0}, \mathbf{I})$, then one obtains the scVI observation model[25]. (We highlight a subtle point: scVI treats the effect of size factors in the measurement model as a random effect whose variance is estimated from the data, where in our discussion we have assumed it is a fixed effect of known magnitude.) Specifically, after marginalizing over $p(u_{ij})$, the decoder network can be written

$$p(\lambda_{ij} \mid l_{i\cdot}) = (\pi(l_{i\cdot}))_j\, \delta_0(\cdot) + \left(1 - (\pi(l_{i\cdot}))_j\right) \text{Gamma}\left(\phi_j^{-1}, (\mu(l_{i\cdot}))_j^{-1} \phi_j^{-1}\right), \tag{32}$$

where $\mu(l_{i\cdot})$ and $\pi(l_{i\cdot})$ are $p$-dimensional neural network outputs and the Gamma distribution is parameterized by shape and rate (i.e., has mean $\mu(l_{i\cdot})_j$ and variance $\mu(l_{i\cdot})_j^2 \phi_j$).

In practice, inference is performed by introducing an *encoder network*[22], which is a neural network $\tilde{f}$ mapping $x_{i\cdot}$ to the approximate posterior distribution of $l_{i\cdot}$ given $x_{i\cdot}$

$$q(l_{i\cdot} \mid x_{i\cdot}) = \text{Normal}(m(x_{i\cdot}), \text{diag}(s^2(x_{i\cdot}))), \tag{33}$$

where $m(\cdot)$ and $s^2(\cdot)$ are $k$-dimensional neural network outputs representing the mean and diagonal of the covariance matrix, and maximizing the evidence lower bound using stochastic gradient descent[22,23]. This algorithm is implemented in the Python package *scvi*. (The software implementation of scVI also supports a Poisson observation model, $u_{ij} = 1$, and an NB observation model, $u_{ij} \sim \text{Gamma}(\phi_j^{-1}, \phi_j^{-1})$.) We note that the description here differs from the original presentation of the method, with consequences on downstream inference (see Supplementary Note 4 for details).

3.1.6. *Poisson variational autoencoder.* If one assumes that $u_{ij} = 1$; that $\mu_{i\cdot} = f(l_{i\cdot})$, where $f$ is parameterized by a neural network; and that $l_{i\cdot} \sim \text{Normal}(\mathbf{0}, \mathbf{I})$, then one obtains the PVAE observation model, which we introduced in the main text. To fit the model, we introduced an encoder network (33). We used two hidden layers of dimension 128 in the encoder and decoder networks, with ReLU activations and batch normalization at each hidden layer. We fit the observation model by maximizing the evidence lower bound using batch gradient descent accelerated by RMSprop[26]. After fitting the observation model, we estimated

$$\hat{\lambda}_{ij} = \frac{1}{S} \sum_{s=1}^{S} (f(l_{i\cdot}^{(s)}))_j, \qquad l_{i\cdot}^{(s)} \sim q(l_{i\cdot} \mid x_{i\cdot}). \tag{34}$$

3.2. **Empirical examples.** In multi-gene models, a common goal is to estimate the underlying low rank structure of the expression matrix $\mathbf{\Lambda}$ (or, in models that include noise, the structured component of expression $\boldsymbol{\mu}$). Our goal here is to address questions such as whether non-linear models provide a better fit to scRNA-seq data than linear models, and whether the use of a specific link function produces a consistently improved fit.

In real data sets, the true value of $\mathbf{\Lambda}$ is unknown. To side-step this problem, we used *binomial thinning*[27,28] of real data sets (independently proposed as *molecular cross-validation*[29]) to produce training and validation data with the same true expression matrix $\mathbf{\Lambda}$. The key idea of this approach is that a Binomial sample of a Poisson-distributed count is also (marginally) Poisson-distributed, meaning it can produce such training/validation data without requiring knowledge of the true $\mathbf{\Lambda}$. We then assessed multi-gene methods by fitting them to the training data, and measuring their predictive accuracy (Poisson log likelihood) on the validation data. The assumption here is that models that more accurately capture the structure in the data should have higher validation log likelihood.

We applied binomial thinning to several data sets and compared three different methods based on different noiseless multi-gene expression models: a linear model assuming identity link (NMF), a linear model assuming log link (GLM-PCA), and a non-linear model (Poisson variational autoencoder, PVAE; Supplementary Note 3). To facilitate comparisons, we implemented these methods within a single Python package *scmodes* (`https://www.github.com/aksarkar/scmodes`), and applied them to sub-samples of 1,000 cells in each data set. We restricted our analysis to genes with non-zero observations in at least 25% of cells (this filter includes all genes in the iPSC data, so we sub-sampled to 500 random genes).

In these comparisons, NMF often performed worse on validation log likelihood than GLM-PCA and PVAE (Supplementary Figure 4a). However, this difference was almost completely explained by behavior in fitting the entries equal to zero in the training data: NMF systematically estimated smaller $\lambda_{ij}$ for these zero entries, leading to poor validation log likelihood in cases where the validation set contained a non-zero count (Supplementary Figure 4b). The Poisson log likelihood is particularly sensitive to small absolute differences (but large relative differences) near 0. Effectively, NMF has a greater tendency than the other methods to overfit the zero entries and underestimate the true expression levels. While this behavior of NMF is statistically undesirable, it could likely be remedied by minor changes (e.g., regularization), and likely does not reflect a fundamental limitation of its assumptions (linear dimension reduction with identity link). Furthermore, it is unclear whether this behavior actually hinders the ability of NMF to capture the important biological structure in the data. Indeed, overall the three methods produce highly concordant estimates of expression $\mathbf{\Lambda}$ (Supplementary Figure 4c).

#### 4. Identifiability of measurement and expression models

Multiple combinations of measurement model and expression model can give rise to the same observation model. Despite the fact that the resulting observation models are the same, and therefore cannot be distinguished from observed data alone, different choices for expression models will naturally give different inferences about true gene expression. We illustrate this fact using the ZINB observation model as an example. The ZINB observation model for counts $x_{1j}, \dots, x_{nj}$ observed at a single gene $j$ is a mixture of two distributions, a point mass at 0 and a Negative Binomial distribution. In our framework, it arises by assuming a point-Gamma expression model (a mixture of a point mass at 0, denoted $\delta_0$, and a Gamma distribution)

$$\begin{aligned} x_{ij} \mid \lambda_{ij}, x_{i+} &\sim \text{Poisson}(x_{i+}\lambda_{ij}), \qquad i = 1, \dots, n \\ \lambda_{ij} \mid \mu_j, \phi_j, \pi_j &\sim \pi_j \delta_0(\cdot) + (1 - \pi_j)\,\text{Gamma}(\phi_j^{-1}, \mu_j^{-1}\phi_j^{-1}). \end{aligned} \tag{35}$$

It is straightforward to show that marginalizing over $\lambda_{ij}$ yields a ZINB distribution on $x_{1j}, \dots, x_{nj}$. In this parametrization, the Gamma component of the point-Gamma distribution has mean $\mu_j$ and variance $\mu_j^2 \phi_j$, and the resulting Negative Binomial component of the ZINB distribution has mean $x_{i+}\mu_j$, dispersion $\phi_j$, and variance $x_{i+}\mu_j + (x_{i+}\mu_j)^2 \phi_j$.

Deriving the ZINB observation model this way clarifies that fitting it to data corresponds to estimating the underlying point-Gamma expression model, removing the effect of measurement error. This point-Gamma expression model contains all of the information about expression variation needed to answer biological questions of interest. For example, under the expression model (35), the mean gene expression is $(1 - \pi_j)\mu_j$.

However, the ZINB observation model can be derived in a different way, using a different measurement model and different expression model, which leads to a different interpretation. Specifically, one could assume a *Zero-inflated Poisson* measurement model (a mixture of a point mass at 0 and a Poisson distribution), and a Gamma expression model

$$\begin{aligned} x_{ij} \mid \lambda_{ij}, x_{i+}, \pi_j &\sim \pi_j \delta_0(\cdot) + (1 - \pi_j)\,\mathrm{Poisson}(x_{i+}\lambda_{ij}) \\ \lambda_{ij} \mid \mu_j, \phi_j &\sim \mathrm{Gamma}(\phi_j^{-1}, \mu_j^{-1}\phi_j^{-1}). \end{aligned} \tag{36}$$

It is straightforward to show that marginalizing over $\lambda_{ij}$ in (36) also leads to a ZINB distribution; in fact, the same ZINB distribution as (35).

Because these two different sets of assumptions, (35) and (36), produce exactly the same observation model, one cannot distinguish between them based on which better fits the observed data better. However, distinguishing between them matters in practice because they lead to different inferences about the actual expression values from the observed data. For example, whereas under (35) the mean of $\lambda_j$ is $(1 - \pi_j)\mu_j$, under (36) it is $\mu_j$. More generally, these two different interpretations could lead to different inferences about e.g., differential expression or clustering[30]. We argue that both theory and empirical evidence support the use of the Poisson measurement model, and not the ZIP measurement model, and that therefore analyses using the ZINB observation model should be derived and interpreted using (35) rather than (36).

#### 5. Negative binomial model for measurement error

To assess whether empirical control data could depart from a Poisson measurement model because of biases in the measurement process that have not been accounted for, we sought to fit an NB measurement model combined with a Gamma expression model at each gene

$$\begin{aligned} x_{ij} \mid x_{i+}, \lambda_{ij}, \theta &\sim \mathrm{NB}(x_{i+}\lambda_{ij}, \theta) \\ \lambda_{ij} \mid \mu_j, \phi_j &\sim \mathrm{Gamma}(\phi_j^{-1}, \mu_j^{-1}\phi_j^{-1}), \qquad j = 1, \dots, p, \end{aligned} \tag{37}$$

where the NB distribution is parameterized by mean and dispersion, and the Gamma distribution is parameterized by shape and rate. Estimating the parameters $\mu_1, \dots, \mu_p, \phi_1, \dots, \phi_p, \theta$ by an EM algorithm requires estimating intractable expectations, and our preliminary results suggest a variational approximation might not be appropriate. To side-step this problem, we note that under (37),

$$\mathrm{E}[x_{ij}] = x_{i+}\mu_j \tag{38}$$

$$\mathrm{V}[x_{ij}] = x_{i+}\mu_j + x_{i+}^2\mu_j^2(\phi_j + \theta + \phi_j\theta), \tag{39}$$

i.e., the first two moments of the compound NB-Gamma observation model equal the corresponding moments of an NB observation model. Using this insight, we approximated the profile likelihood of the data with respect to $\theta$. For each candidate measurement dispersion level $\theta$, we fit an NB observation model (Poisson measurement model combined with a Gamma expression model) by maximizing the likelihood, yielding an estimate of $d_j = \phi_j + \theta + \phi_j \theta$. If the estimate $\hat{d_j}$ implies $\phi_j \leq 0$, then we fit the Gamma expression model fixing $d_j = 1/\theta$.

#### 6. Supplementary Methods

6.1. **Empirical analysis of single-gene expression models.** To assess questions such as whether a Gamma expression model is adequate to model expression variation at most genes, or whether it is necessary to use a more complex, even multi-modal, expression model, we compared different expression models by estimating the likelihood of observed data under each model. Specifically, we compared a Gamma distribution, a point-Gamma distribution, a non-parametric unimodal distribution, and a fully non-parametric distribution. Since these comparisons involve non-parametric families, and specifying any of these models as a null expression model (to obtain $p$-values) is also questionable, we considered a likelihood ratio of 100 or more as strong evidence for one model over another as a simple heuristic.

We analyzed each gene in several data sets from a range of settings including homogeneous collections of sorted cells, *a priori* homogeneous cell lines, and heterogeneous tissues (Table 3). We also created *in silico* mixtures of sorted cells as positive controls for highly heterogeneous expression patterns. We downloaded data from Gene Expression Omnibus or the Human Cell Atlas for previously published data, and directly from 10X Genomics for Chromium v3 data. We did not include observed covariates in the observation model, since for most data sets these were not available.

To fit Gamma expression models of the form

$$\lambda_{ij} \sim \operatorname{Gamma}(a_j, b_j), \tag{40}$$

and point-Gamma expression models of the form

$$\lambda_{ij} \sim \pi_j \delta_0(\cdot) + (1 - \pi_j) \operatorname{Gamma}(a_j, b_j), \tag{41}$$

where the Gamma distribution is parameterized by shape and rate, we used an EM algorithm[8] accelerated by SQUAREM[31], implemented in the Python package *scmodes* (https://www.github.com/aksarkar/scmodes).

To fit non-parametric unimodal expression models, we used generalized adaptive shrinkage, implemented in the R package *ashr*. These expression models take the form

$$\lambda_{ij} \sim \sum_{k=1}^{K_j} w_{jk} \operatorname{Uniform}(\cdot; \lambda_{0j}, a_{jk}), \tag{42}$$

where the number of endpoints $K_j$ and their locations $a_{jk}$ are fixed based on the range of the data, and $\operatorname{Uniform}(\cdot; \lambda_{0j}, a_{jk})$ denotes the density of the uniform distribution on $[a_{jk}, \lambda_{0j}]$ if $a_{jk} < \lambda_{0j}$, and on $[\lambda_{0j}, a_{jk}]$ if $a_{jk} > \lambda_{0j}$.

To fit fully non-parametric expression models, we used generalized adaptive shrinkage (*ashr*) as a subroutine in an iterative procedure, implemented in the Python package *scmodes*. These expression models take the form,

$$\lambda_{ij} \sim \sum_{k=1}^{K_j} w_{jk} \operatorname{Uniform}(\cdot; (k-1)a_j, ka_j), \tag{43}$$

where the number of components $K_j$ is fixed and the grid step size $a_j$ depends on the range of the data. We initially fit an expression model with $K_j = 512$, and in each iteration discarded mixture components with estimated $w_{jk} < 10^{-8}$ and divided each remaining mixture component $k$ into two components by dividing the segment $[(k-1)a_j, ka_j]$, and the associated mixture weight $w_{jk}$, in half. We terminated this procedure when the change in log likelihood was less than $10^{-7}$, or after 40 iterations.

6.2. **Empirical analysis of alternative measurement models.** To bound the level of measurement dispersion that is consistent with observed data, we made an assumption that the level of overdispersion in the measurement process is the same for all genes. Under this assumption, it is possible to estimate the measurement dispersion by fitting an NB observation model with separate dispersion parameters to reflect measurement dispersion $\theta$ and uncontrolled expression variation $\phi_j$

$$\begin{aligned} x_{ij} \mid x_{i+}, \lambda_{ij} &\sim \text{Poisson}(x_{i+}\lambda_{ij}) \\ \lambda_{ij} \mid \mu_j, d_j &\sim \text{Gamma}(d_j^{-1}, \mu_j^{-1} d_j^{-1}), \end{aligned} \tag{44}$$

where $d_j \triangleq \phi_j + \theta + \phi_j\theta$ and the Gamma distribution is parameterized by shape and rate (see Supplementary Note 5 for details). We estimated the profile log likelihood of the data under this model $L(\theta)$ as a function of $\theta$ by fitting the model (44) over a grid of candidate values for $\theta$. For a fixed value of $\theta$, if the estimate $\hat{d}_j$ implies $\phi_j \leq 0$, then we fit the observation model (44) fixing $d_j = 1/\theta$. We implemented this procedure in the Python package *scmodes*.

We applied this approach to five control data sets spanning four experimental protocols which were previously pre-processed and analyzed[3,32] (Table 3). To determine the bound, we chose the maximum level of measurement dispersion $\theta$ for which $L(\theta)/L(\theta^*) \geq 0.1$, where $\theta^*$ is the maximum likelihood estimate.

6.3. **Empirical analysis of multi-gene expression models.** To assess the accuracy of methods in estimating the true gene expression levels $\mathbf{\Lambda}$, we used binomial thinning[27,28] of real data sets (independently proposed as molecular cross-validation[29]) to produce training and validation data with the same true expression matrix $\mathbf{\Lambda}$. We then assessed methods by fitting them to the training data, and estimating the log likelihood of the validation data. The assumption here is that models that more accurately capture the structure in the data should have higher validation log likelihood.

We compared three different noiseless multi-gene expression models: a linear model assuming identity link (Poisson NMF[14–17]), a linear model assuming log link (GLM-PCA[18]), and a non-linear model (Poisson variational autoencoder, PVAE; see Supplementary Note 3 for details). To facilitate comparisons, we implemented these methods in the Python package *scmodes*.

We applied each method to several biological data sets (Table 3). In each data set, we took 10 independent, random sub-samples of 1,000 cells. We restricted our analysis to genes with non-zero observations in at least 25% of cells. This filter includes all genes in the iPSC data set, so we additionally sub-sampled this data set to 500 random genes.

#### 7. Supplementary Figures

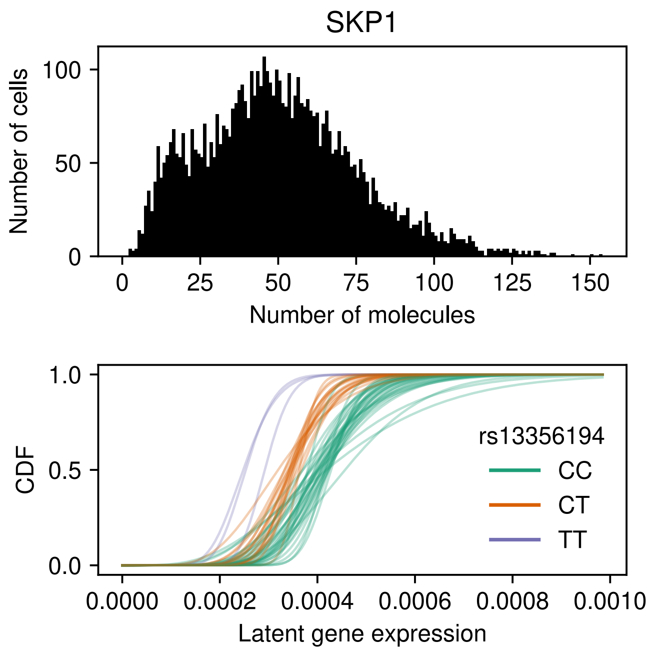


Supplementary Figure 1. **Structured expression variation at *SKP1* in iPSCs.** Example gene in iPSCs that shows weak evidence for a fully non-parametric expression model over a unimodal model when analyzing all samples jointly, but exhibits multimodal variation when analyzed separately by donor. Color indicates genotype at a nearby eQTL.

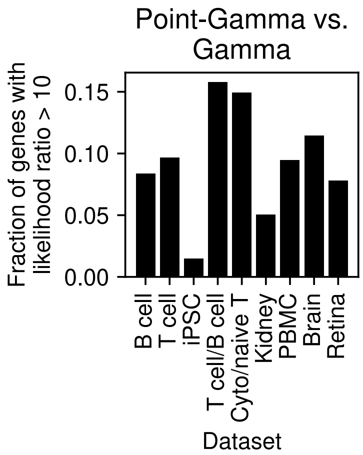


Supplementary Figure 2. **Evidence for an excess of zeros in biological data sets.** Fraction of genes in biological data sets with suggestive evidence in favor of a point-Gamma expression model against a Gamma expression model.

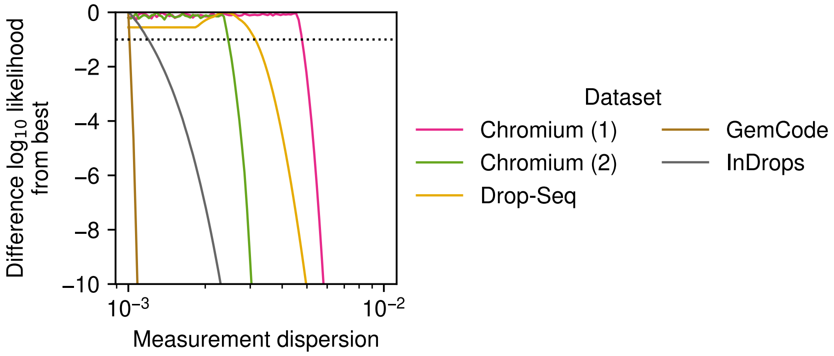


Supplementary Figure 3. **Evidence for an overdispersed measurement model.** Profile log likelihood of control data as a function of the level of measurement dispersion in an NB measurement model.

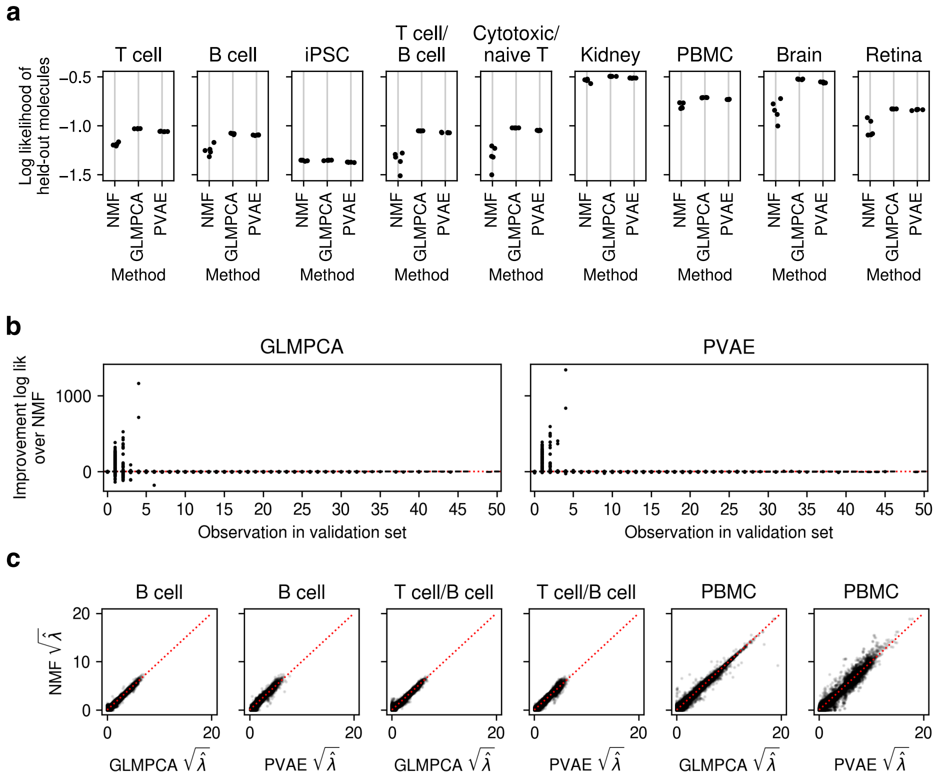


Supplementary Figure 4. **Goodness of fit of multi-gene models of scRNA-seq.** (a) Per observation (sample-gene combination) log likelihood of held out molecules for 5 random training/validation splits, fixing the rank of each model to 8. (b) Distribution of log likelihood differences on one validation data set generated by holding out molecules from the B cells data set. (c) Comparison of estimated $\hat{\boldsymbol{\Lambda}}$ for three representative training data sets, fixing the rank of each model to 8.

### References

1. Baker, S. G. The Multinomial-Poisson Transformation. *J Royal Stat Soc* **43,** 495–504 (1994).
2. Pachter, L. Models for transcript quantification from RNA-Seq. *arXiv e-prints* (2011).
3. Wang, J. *et al.* Gene expression distribution deconvolution in single-cell RNA sequencing. *Proc Natl Acad Sci USA* (2018).
4. Sarkar, A. K. *et al.* Discovery and characterization of variance QTLs in human induced pluripotent stem cells. *PLoS Genetics* **15,** 1–16 (2019).
5. Zeileis, A., Kleiber, C. & Jackman, S. Regression Models for Count Data in R. *Journal of Statistical Software* **27** (2008).
6. Venables, W. N. & Ripley, B. D. *Modern Applied Statistics with S* Fourth. ISBN 0-387-95457-0 (Springer, New York, 2002).
7. Adamidis, K. Theory & Methods: An EM algorithm for estimating negative binomial parameters. *Australian & New Zealand Journal of Statistics* **41,** 213–221 (1999).
8. Karlis, D. EM Algorithm for Mixed Poisson and Other Discrete Distributions. *ASTIN Bulletin* **35,** 3–24 (2005).
9. Stephens, M. False discovery rates: a new deal. *Biostatistics* **18,** 275–294 (2017).
10. Lu, M. *Generalized Adaptive Shrinkage Methods and Applications in Genomics Studies* PhD thesis (University of Chicago, 2018).
11. Efron, B. Empirical Bayes deconvolution estimates. *Biometrika* **103,** 1–20 (2016).
12. Kiefer, J. & Wolfowitz, J. Consistency of the Maximum Likelihood Estimator in the Presence of Infinitely Many Incidental Parameters. *Ann Math Statist* **27,** 887–906 (1956).
13. Koenker, R. & Mizera, I. Convex Optimization, Shape Constraints, Compound Decisions, and Empirical Bayes Rules. *J Am Stat Assoc* **109,** 674–685 (2014).
14. Zhu, X., Ching, T., Pan, X., Weissman, S. & Garmire, L. Detecting heterogeneity in single-cell RNA-Seq data by non-negative matrix factorization. *PeerJ,* 5:e2888 (2017).
15. Matsumoto, H. *et al.* An NMF-based approach to discover overlooked differentially expressed gene regions from single-cell RNA-seq data. *NAR Genomics and Bioinformatics* **2.** lqz020 (2019).
16. Lee, D. D. & Seung, H. S. *Algorithms for Non-negative Matrix Factorization* in *Advances in Neural Information Processing Systems 13, Papers from Neural Information Processing Systems (NIPS) 2000, Denver, CO, USA* (eds Leen, T. K., Dietterich, T. G. & Tresp, V.) (MIT Press, 2000), 556–562.
17. Cemgil, A. T. Bayesian Inference for Nonnegative Matrix Factorisation Models. *Comput Intell Neurosci* (2009).
18. Townes, F. W., Hicks, S. C., Aryee, M. J. & Irizarry, R. A. Feature selection and dimension reduction for single-cell RNA-Seq based on a multinomial model. *Genome Biol* **20,** 295 (2019).
19. Risso, D., Perraudeau, F., Gribkova, S., Dudoit, S. & Vert, J.-P. A general and flexible method for signal extraction from single-cell RNA-seq data. *Nat Commun* **9,** 284 (2018).
20. Eraslan, G., Simon, L. M., Mircea, M., Mueller, N. S. & Theis, F. J. Single-cell RNA-seq denoising using a deep count autoencoder. *Nat Commun* **10,** 390 (2019).
21. Dayan, P. in *The Handbook of Brain Theory and Neural Networks* 522–524 (MIT Press, 2003).

22. Kingma, D. P. & Welling, M. *Auto-Encoding Variational Bayes* in *2nd International Conference on Learning Representations, ICLR 2014, Banff, AB, Canada, April 14-16, 2014, Conference Track Proceedings* (eds Bengio, Y. & LeCun, Y.) (2014).
23. Rezende, D. J., Mohamed, S. & Wierstra, D. *Stochastic Backpropagation and Approximate Inference in Deep Generative Models* in *Proceedings of the 31st International Conference on Machine Learning* (eds Xing, E. P. & Jebara, T.) **32** (PMLR, Bejing, China, 2014), 1278–1286.
24. Gershman, S. & Goodman, N. *Amortized inference in probabilistic reasoning* in *Proceedings of the annual meeting of the cognitive science society* **36** (2014).
25. Lopez, R., Regier, J., Cole, M. B., Jordan, M. I. & Yosef, N. Deep generative modeling for single-cell transcriptomics. *Nat Methods* **15,** 1053–1058 (2018).
26. Tieleman, T. & Hinton, G. *Lecture 6.5—RmsProp: Divide the gradient by a running average of its recent magnitude* COURSERA: Neural Networks for Machine Learning. 2012.
27. Gerard, D. & Stephens, M. Unifying and Generalizing Methods for Removing Unwanted Variation Based on Negative Controls. *arXiv e-prints* (2017).
28. Gerard, D. Data-based RNA-seq simulations by binomial thinning. *BMC Bioinformatics* **21,** 206 (2020).
29. Batson, J., Royer, L. & Webber, J. Molecular Cross-Validation for Single-Cell RNA-seq. *bioRxiv* (2019).
30. Silverman, J. D., Roche, K., Mukherjee, S. & David, L. A. Naught all zeros in sequence count data are the same. *bioRxiv* (2018).
31. Varadhan, R. & Roland, C. Simple and Globally Convergent Methods for Accelerating the Convergence of Any EM Algorithm. *Scandinavian Journal of Statistics* **35,** 335–353 (2008).
32. Svensson, V. Droplet scRNA-seq is not zero-inflated. *Nat Biotech* (2020).
